# Validating Antibodies for Quantitative Western Blot Measurements with Microwestern Array

**DOI:** 10.1101/221523

**Authors:** Rick J. Koch, Anne Marie Barrette, Alan D. Stern, Bin Hu, Evren U. Azeloglu, Ravi Iyengar, Marc R. Birtwistle

**Affiliations:** Department of Pharmacological Sciences, Icahn School of Medicine at Mount Sinai, New York, NY 10029, USA; Department of Chemical and Biomolecular Engineering, Clemson University, Clemson, SC 29634, USA

## Abstract

Western blotting is often considered a semi-quantitative or even qualitative assay for assessing changes in protein or protein post-translational modification levels. Fluorescence-based measurement enables acquisition of quantitative data in principal, but requires determining the linear range of detection for each antibody—a labor-intensive task. Here, we describe the use of a high-throughput western blotting technique called microwestern array to more rapidly evaluate suitable conditions for quantitative western blotting with particular antibodies. We can evaluate up to 192 antibody/dilution/replicate combinations on a single standard size gel with a seven-point, two-fold lysate dilution series (~100-fold range). Pilot experiments demonstrate a surprisingly high proportion of investigated antibodies (17/22) are suitable for quantitative use, and that lack of validity might often be a consequence of lysate composition rather than antibody quality. Linear range for all validated antibodies is at least 8-fold, and in some cases nearly two orders of magnitude. That range could be greater as the presented tests did not find a limit for many antibodies. We find that phospho-specific and total antibodies do not have discernable trend differences in linear range or limit of detection, but total antibodies generally required higher working concentrations, suggesting phospho-specific antibodies may be generally higher affinity. Importantly, we demonstrate that results from microwestern analyses scale to normal “macro” western for a subset of antibodies. These data indicate that with initial validation, many antibodies can be readily used quantitatively in a reproducible manner. Antibody validation data and standard operating procedures are available online (www.birtwistlelab.com/protocols and www.dtoxs.org).

## INTRODUCTION

Scientific research, and in particular that in the biomedical field, has come under harsh scrutiny and debate of late due to questions of reproducibility [1–10]. While there are many potential reasons for lack of reproducibility, one major reason relates to research reagents, including antibodies [11–13]. Antibodies are widely-used critical tools in a variety of biomedical research assays, but they are not always suitable for the application of interest. The intended application for the antibody brings potentially different criteria and stringency for their use. For example, qualitative inference from immunohistochemistry may be possible, but acquiring quantitative data from flow cytometry may not be with the same antibody and cell system. Antibody validity is highly dependent on biological context and the assay itself[11–13].

One major application of antibodies in both large and small labs is the western blot. While the western blot is often considered semi-quantitative or qualitative, it can be quantitative with infrared fluorescence-based detection[14–18]. Reverse phase protein array (RPPA) is a well-established method for quantitative data from cell and tissue lysates[19–22], but it does not separate proteins by molecular weight, and therefore has more stringent requirements for antibody validity. In fact, RPPA protocols report using western blotting as the method for validating antibodies for RPPA use [22].

Here, we focus on showing how a meso-scale western blotting platform called microwestern array can help provide information to assess the validity of quantitative data from western blots—a form of antibody validation. This of course considers that other important aspects of antibody validation, such as specificity via genetic approaches, are already validated[12]. The microwestern array was originally developed in 2011 in the Jones lab at the University of Chicago[23,24]. The microwestern process is very similar to the regular “macro” western; lysates are in an SDS-containing buffer and proteins are separated by molecular weight via electrophoresis, transferred to a membrane, and incubated with antibodies for detection (Fig. 1). The major difference is that lysates are spotted onto the surface of a gel via piezo-electric pipetting, which allows for incubation with up to 192 antibodies (96x2 colors) via a gasketed hybridization plate after transfer to the membrane. We have implemented the microwestern array in the context of our NIH Library of Integrated Network Cellular Signatures (LINCS) Data Generation Center[25–27]. One major thrust of LINCS is improving data FAIR-ness (Findable, Accessible, Interoperable, Reusable)[28], and this particular application of microwestern is one aspect of LINCS focusing on reagent validation that is critical in such endeavors. We show that results from microwestern scale to regular western. We provide an initial set of data that investigates such validity across a set of evaluated antibodies which will continue to grow and be publicly available.

**Figure 1.**
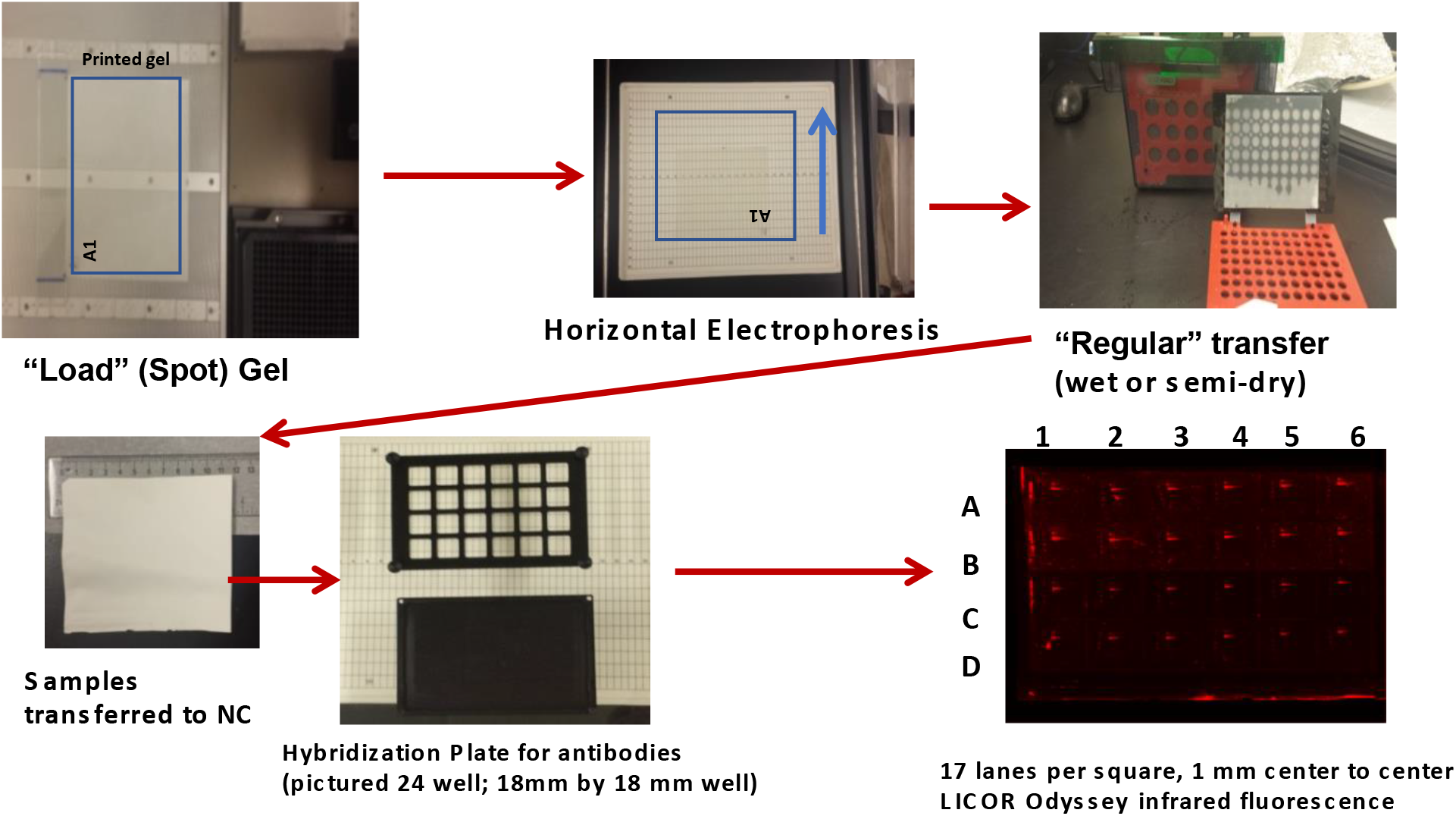
Work Flow for the Microwestern. The major steps involved in the Microwestern protocol from spotting the samples onto the gel to the final image showing sample signals in a 24 well setup. Details are presented in Methods and the referenced SOPs therein.

## METHODS

### Reagents and Sources

For the below, detailed protocols and standard operating procedures can also be found on the Birtwistle Lab website (www.birtwistlelab.com/protocols) or the DToxS website (www.dtoxs.org). These protocols contain detailed reagent and source information where not listed explicitly below.

### Cell Culture

MCF10A cells were plated at 2 million cells per 100mm dish, cultured overnight in serum containing media and harvested approximately 24 hours later. Media was aspirated and cells were prepared for lysis by washing 2x in ice-cold phosphate-buffered saline (PBS). PromoCells were cultured as described in SOP_CE-2.0-PromoCell_Myocyte_Plating_for_Drug_Test on www.dtoxs.org.

### Collection of Total Protein for Ab Validation

Primary human cardiac myocytes (PromoCells) and MCF10A cells were lysed using an aqueous buffer consisting of 240mM Tris-acetate, 1% w/v SDS, 0.5% v/v glycerol, 5mM EDTA, to which the following inhibitors were added immediately prior to use; aprotinin, leupeptin, pepstatin, beta galactophosphatase, activated sodium orthovanadate and DTT. 1.0 mL ice-cold lysis buffer was added per 10 cm dish after washing as above. Cells were collected with a cell scraper and lysate transferred into a pre-cooled Eppendorf Protein LoBind 1.5mL tube using a micropipette. Volumes of lysate consisted of 1ml or more total volume which were divided into approximately 500ul aliquots and placed into LoBind tubes on ice. Lysates were sonicated using the Hielscher Ultrasonics VialTweeter placed in a 4°C cold room using 10 cycles, each cycle consisting of 30 seconds sonication at 100% amplitude followed by a 30 second rest period. Tubes were then placed into a 95°C heat block for 2 minutes and stored at −80°C until processed. Tubes were then placed on ice to thaw and heated at 95°C in a heat block for 2 minutes then immediately aliquoted into Amicon 500uL Centricon spin columns for concentration. Lysates were spun at 14,000g, room temperature for 15min. Each aliquot was concentrated ~5X to 10X by volume. Concentrated aliquots were pooled and 4uL was set aside for total protein measurement by Pierce 660 with the remainder frozen at −80°C until use. The aliquot used in the Pierce 660 protein assay was diluted 10 to 20 fold with lysis buffer to bring it to a concentration of about 1mg/ml, within the range of the BSA standards used. Based on the 660 results, lysate was diluted with lysis buffer to produce the 2-fold dilution series used in Ab validation.

### Printing of Samples onto Micro western Gel

The lysate samples ranging from 10mg/ml to 0.31mg/ml (predominantly, with some exceptions as noted) along with MW standards were printed onto a 9.5% acrylamide gel cast especially for a Microwestern (see SOP# A 9.0 Casting of Gel on www.dtoxs.org) using the GeSim Nanoplotter Model 2.1E with the GeSim software NPC16V2.15.53. The appropriate workplate file was loaded depending upon the configuration of the hybridization plate used, either the 24 or 96 well plate (available upon request). Also, according to the type of hybridization plate used and the exact samples printed, a unique Transfer file was run to load samples, appropriately aliquoted into a 384-well black microtiter plate, onto the gel. The Transfer file used printed approximately 75 nL of each sample; ~500 pL per spot, 15 spots per cycle, 10 cycles. (See SOP# A11.0, Printing Gel; www.dtoxs.org).

### Electrophoresis and Transfer to Nitrocellulose

Printed samples were electrophoresed either about 8mm or 15mm according to the hybridization plate used (96 or 24 well, respectively) with a horizontal electrophoretic box (Gel Company) prechilled to 10°C with a Huber Minichiller. (See SOP# A 13.0, Microwestern Electrophoresis, www.dtoxs.org). The gel containing the MW-separated samples was then carefully placed onto a transfer buffer (Tris/Glycine/MeOH—see SOP# A 14.0, Microwestern Wet Transfer) dampened filter paper, samples facing up, so that the sample region overlaid the filter paper. The filter paper is part of the BioRad filter paper/nitrocellulose/filter paper sandwich. The NC was wetted with transfer buffer and precisely laid on top of the gel without movement of the NC after placement. Any air bubbles were carefully rolled out, the remaining filter paper wetted with transfer buffer was placed onto the NC, and the sandwich was placed into the blotter gel holder cassette and samples were transferred overnight at 4°C. (See SOP# A 14.0, Microwestern Wet Transfer, www.dtoxs.org).

### Antibody Incubation, Membrane Imaging and Quantification

Nitrocellulose containing samples was removed from the transfer cassette and placed in Odyssey blocking buffer for 30 minutes to 1 hour at room temperature. Primary antibodies were prepared according to the type of hybridization plate used and the Ab concentration tested. The 24 well plate and 96 well plate use 500uL and 100uL per well, respectively. Antibodies were diluted with the Odyssey blocking buffer. Nitrocellulose was removed from blocking buffer and trimmed for proper alignment within the hybridization plate. Sample side up NC was carefully aligned so printed areas matched the location of the hybridization wells. Antibodies were added into wells using an antibody plate map to ensure the appropriate antibody – well mapping. The plate was covered with optical adhesive film and incubated with gentle rocking overnight at 4°C. Ambient light was blocked with aluminum foil. Primary antibody was aspirated from the plate and NC was washed 5x for about 5 minutes each with TBST (20mM Tris; 137mM NaCl; 0.1% Tween) while still in the plate with washing buffer, while protected from ambient light. Goat anti-mouse and/or goat anti-rabbit secondary antibodies were prepared in the appropriate buffer. After washing (as above), the NC was incubated with secondary antibody for 1 hour at room temperature. Nitrocellulose was again washed and then allowed to dry at room temperature. During all wash steps and drying NC was protected from light. Putting sample side down the NC was scanned using the Li-Cor Odyssey Clx scanner set at “Auto.” (See SOP# A 15.0, “Antibody Incubation,” www.dtoxs.org). Signal was quantified using the Li-Cor “Image Studio Lite, Ver. 5.2 software. (See SOP A 16.0, “Quantifying the Image for Microwestern Array,” www.dtoxs.org)

## RESULTS AND DISCUSSION

### Repeatability of Piezo-Electric Pipetting Sets Upper Bounds for Performance

The microwestern uses a piezo-electric pipetting apparatus to spot lysate onto a gel. The repeatability of this pipetting apparatus will therefore set an upper limit to the quantitative performance of an antibody as evaluated by microwestern. To assess this repeatability, we arrayed a two-fold serial dilution series of molecular weight (MW) ladder onto each well of a 24-well layout, performed electrophoresis, and transferred to nitrocellulose for imaging (Fig. 2A). The MW ladder is fluorescent and therefore quantifiable. Two example quantification results are shown, one near the best performance (Fig. 2B), and one near the worst performance (Fig. 2C). We evaluate performance by the R^2^ value between the amount of MW ladder spotted and the quantified fluorescence intensity (using the 50 kDa spot as a representative proxy), taken as an average across at least three replicate wells. These results demonstrate we can expect the upper range of performance for R^2^ to be between 0.97 and 0.99. We use these criteria to place results for antibody relationships in an appropriate context. Namely, above an R^2^ of 0.97, an antibody may be interpreted to have suitable quantitative behavior. In this paper we consider down to 0.95 as potentially acceptable, although such decisions are of course at the discretion of the researcher.

**Figure 2.**
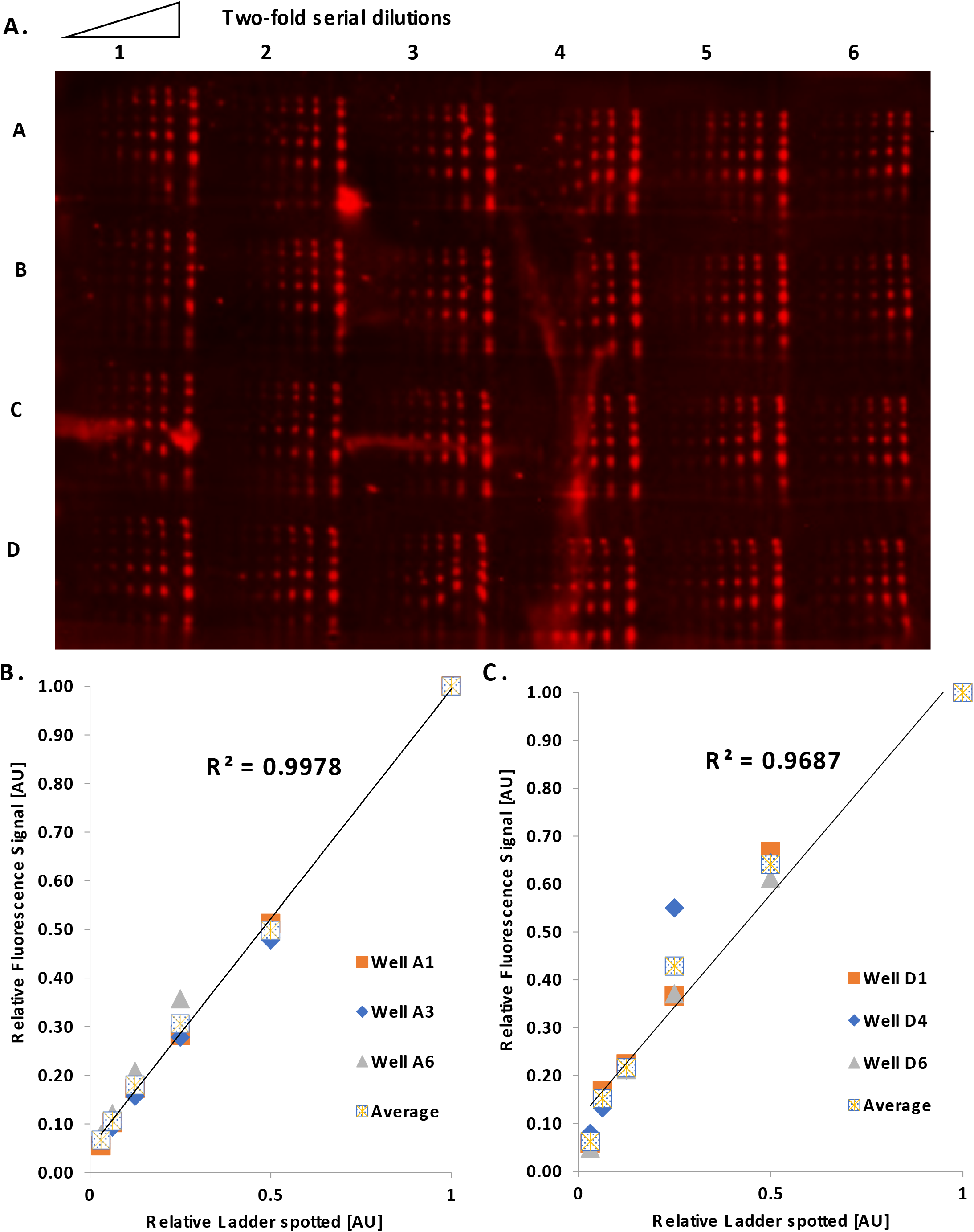
Repeatability of the Piezo Electric Pipetting Device. Using a molecular weight ladder diluted from 1/2 strength to 1/64 in 2-fold steps using lysis buffer to determine the upper most and lowest acceptable values for Ab validation. The 50 kDa band is used for quantification. A. Scan of 24 well setup used to measure signal of molecular weight ladder. B. Analysis of 3 random wells in Row A establishing high R^2^. C. Analysis of 3 random wells in Row D establishing low R^2^.

### Evaluating Quantitative Antibody Performance

To demonstrate a typical antibody validation, we focus on data for doubly phosphorylated ERK1/2 (ppERK-Fig. 3). Lysates from exponentially growing MCF10A cells were spotted in two-fold serial dilutions across 5 points, from 6 mg/mL to 0.37 mg/mL. Six wells of a 24-well setup are shown that corresponded to triplicates with two different primary antibody dilutions (1:2000 and 1:4000). A single spot at the expected molecular weight is observed (representative image in Fig. 3B). Images were quantified, and the averages across triplicates were plotted versus the relative amount of spotted lysate to evaluate R^2^. Both primary antibody dilutions yielded excellent R^2^ values (Fig. 3C). We therefore deemed this antibody as suitable for quantitative use, and recommend the lowest tested primary dilution (1:4000).

**Figure 3.**
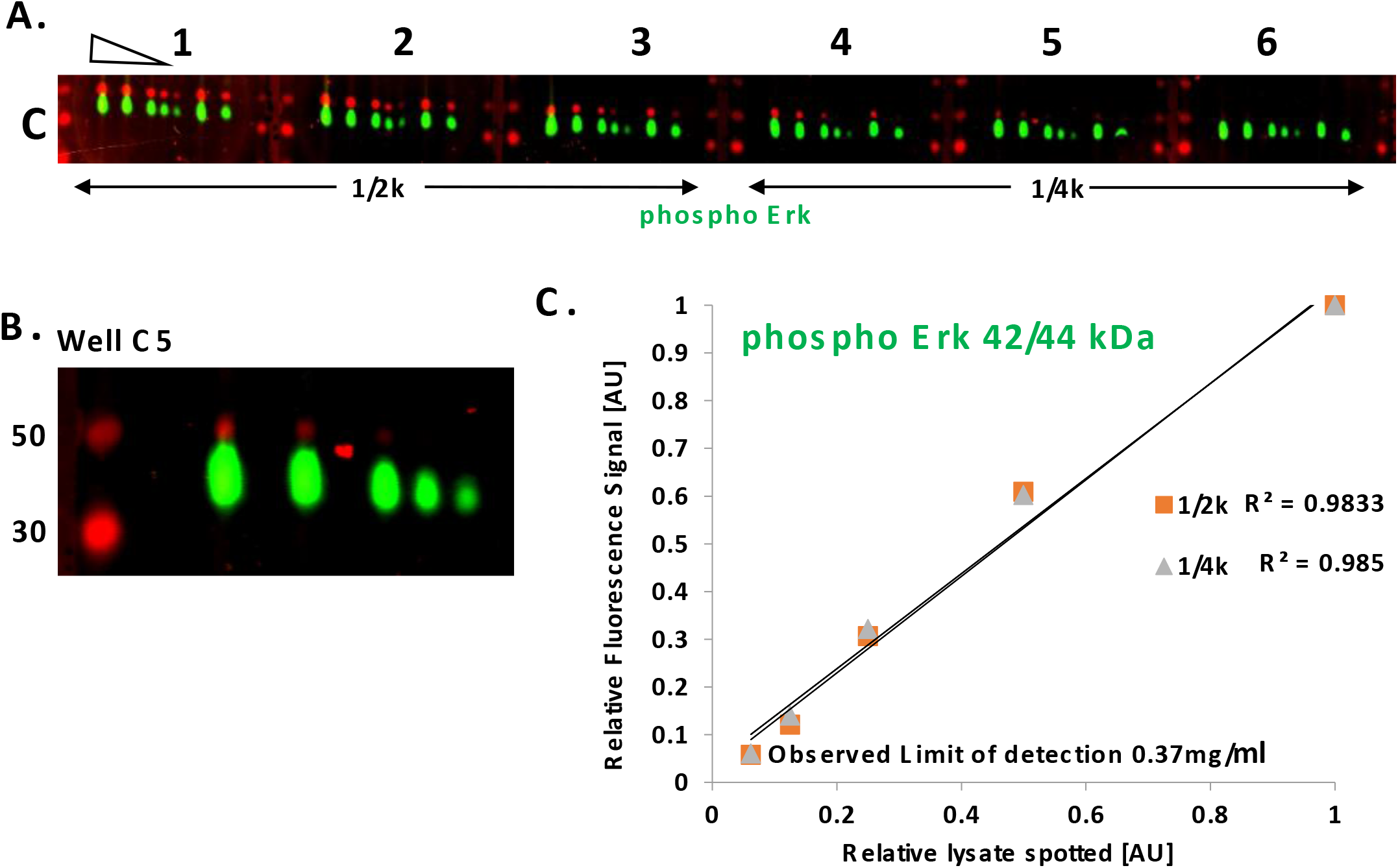
Example of Ab Validation Data. A 5 point 2-fold dilution series of MCF10A cell lysate was used to measure signal for different primary ppERK1/2 antibody concentrations (2 are shown). A. Wells C1 – C6 of a 24 well setup used to measure ppERK signal at 1/2000 and 1/4000 antibody dilution. B. Well C5 showing signal at the correct M.W. of 42/44kDa. C. Analysis of signal from each group of triplicates for each of the antibody dilutions.

This particular 24-well MWA run included data to evaluate other antibodies. We have run other such antibody validation runs in a 96-well format, which allows for denser antibody testing in each gel (Fig. 4A). With so many antibodies being tested from one gel, all spanning large ranges of signal intensity, it is understandably not possible to capture ideal image display across them all. We have additionally taken advantage of the fact that each well can include two different antibodies, so long as they are from different species (e.g. mouse and rabbit), since we image two separable infrared colors. The compilation of our antibodies validated for quantitative use thus far are shown in Table 1. We have validated 17 out of 22 tested. The other five tested but not validated are shown in Table 2.

**Figure 4.**
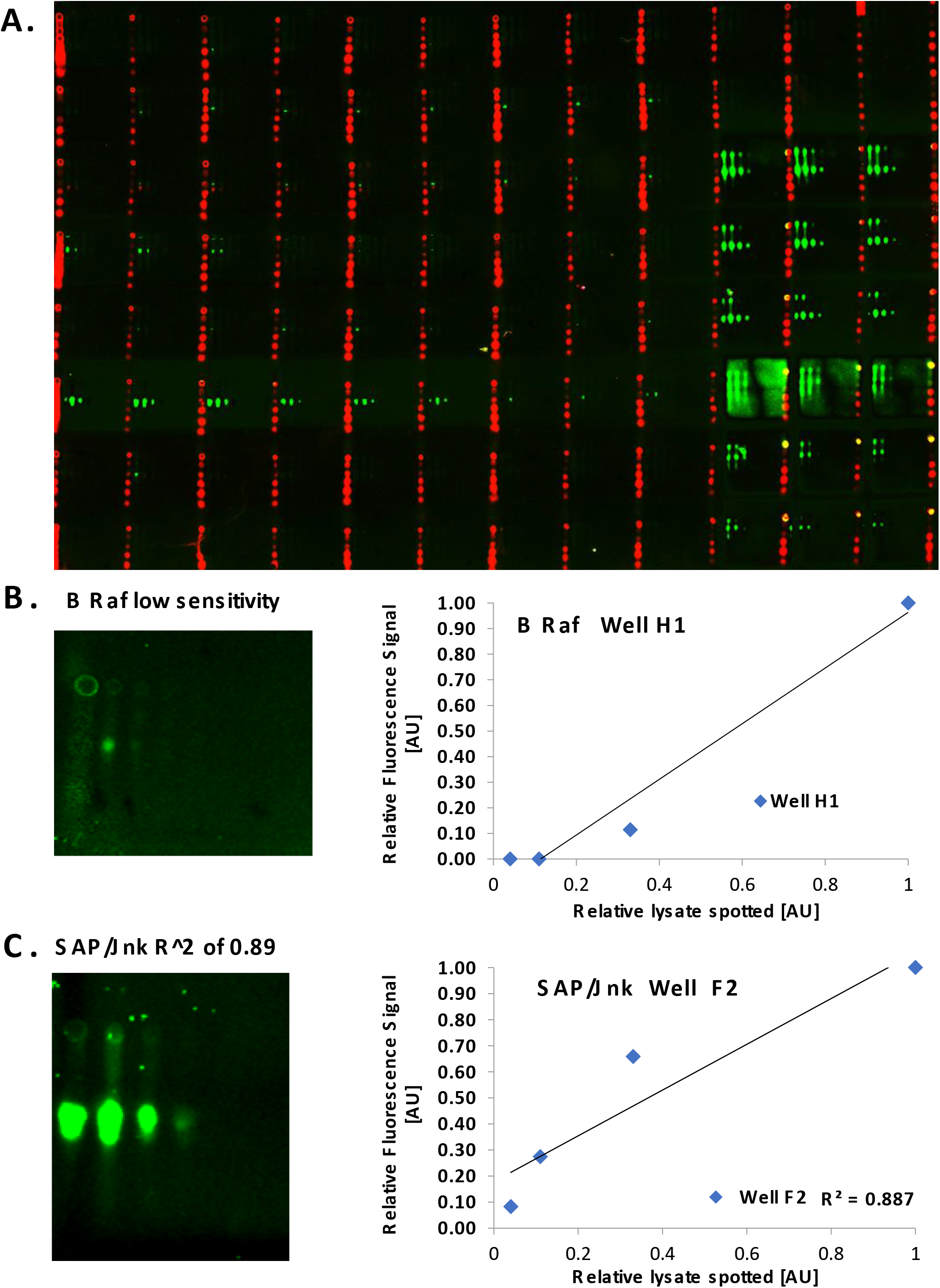
Alternative Ab Validation Format and Failed Validation. A. Representative image from a 96 well setup used for Ab validation. B. Example of failed validation due to low sensitivity, failed to detect 1.11 mg/ml and 0.37 mg/ml lysate samples. C. Example of failed validation due to technical issues. Signal values giving a very low R^2^ value of 0.89.

**Table 2.** Antibodies Not Validated

| Antibody | Mouse/<br>Rabbit | Company | Catalog # | Ref/Lot# | Cell type | Treatment | Total protein -<br>highest<br>mg/ml<br>tested | Range and [Ab]<br>tested |  | Invalid Due to |  |  |
| --- | --- | --- | --- | --- | --- | --- | --- | --- | --- | --- | --- | --- |
|  |  |  |  |  |  |  |  | Total<br>protein<br>mg/ml | [Ab] | Sensitivity | Linearity | Wrong<br>M.W./Multiple<br>bands |
| PI 3 Kinase p85<br>(19H8) | rabbit | Cell<br>Signaling | CS#4257P | 04/2015 | MCF10A | w/serum | 10 | 10 - 0.31 | 1/250;<br>1/500;<br>1/1k | X |  |  |
| cRaf | mouse | Cell<br>Signaling | CS#12552S | 06/2015 | MCF10A | w/serum | 10 | 10 - 0.31 | 1/250;<br>1/500;<br>1/1k | X |  |  |
| CAM | mouse | Cell<br>Signaling | CS#50049S | 06/2015 | MCF10A | w/serum | 10 | 10 - 0.31 | 1/250;<br>1/500;<br>1/1k | X |  |  |
| GSK 3b (3D10) | mouse | Cell<br>Signaling | CS#9832S | 07/2015 | MCF10A | w/serum | 10 | 10-0.37 | 1/500;<br>1/1k;<br>1/2k | X |  |  |
| B - Raf (55C6) | rabbit | Cell<br>Signaling | CS#9433 | 07/2015 | MCF10A | w/serum | 10 | 10-0.37 | 1/500;<br>1/1k;<br>1/2k | X |  |  |

### Primary Antibody Dilutions and Differences in Cell Context

We investigated primary antibody dilutions from 1/250 to 1/4000 across our tests, and we found adequate dilutions at both those extremes, as well as in-between. A typical starting point for western blotting is 1/1000, and this was very frequently an adequate dilution. When not 1/1000, most often the recommended dilution was lower, at 1/250 or 1/500. These dilutions are seldom used in western blotting, but may actually be required for rigorous quantitative analysis in many cases. Only three antibodies have recommended dilutions greater than 1/1000, and notably, one of those (Akt pan) was recommended at 1/250 in a different cell context. This highlights the fact that it is possible that in different cell systems or treatment conditions, the amount of epitope can be different, so the reported primary antibody dilution may be considered a rule of thumb when applied to other such contexts. We found this to be the case in all such examples where we tested the same antibody in those two different cell contexts (ERK1/2, P38MAPK, and phospho-GSK3beta). We noted that in the cases with the largest antibody recommended dilution differences across cell type, it expectedly trended with limit of detection. That is, lower limit of detection implies more primary antibody is needed for rigorous quantitation.

### Properties of Linear Range and Limit of Detection

Besides showing acceptable R^2^, we evaluated linear range of detection as well as the limit of detection for each antibody (Table 1). Antibodies were quantitatively valid between 8-fold and 64-fold ranges—with most at 16-fold. This range captures most biologically relevant changes in total or phospho-protein levels. Limit of detection was typically in the range of 0.2–0.4 mg/mL of total protein in the lysate, with a few higher exceptions. We expect this to be an upper bound when comparing to macro-western, due to the small sample size and sample loss during adsorption onto a gel (versus embedding into wells). Furthermore, the limit of detection and linear range often spanned the edges of that tested. We did not pursue experiments specifically dedicated to expanding these ranges, so the reported properties may be regarded as conservative in this way as well.

### Comparing Antibodies Against Total and Phospho-Protein Epitopes

We wondered whether we could detect systematic differences between phospho-specific antibodies (6 tested), and those against total protein (16 remaining). We first noted that all five antibodies that could not be validated were against total protein. Notably, two of these (c-Raf and GSK3beta) had phospho-specific antibodies that were validated. Moreover, phospho-specific antibodies tended to require lower working concentrations, and did include the two cases where 1/4000 dilutions were validated (phospho-ERK1/2 and phospho-Akt). Notably, the antibodies against the cognate total proteins (ERK1/2 and Akt-pan) required much higher working concentrations. These observations suggest that phospho-antibodies may generally have higher affinity than those against total protein, although definitive conclusions are precluded based on our sample size thus far. In contrast, we could not detect any differences with respect to linear range or limit of detection.

### Reasons for Failure

It is useful to analyze reasons why antibodies fail validation for quantitative use. Fig. 4B-C show two such reasons we encountered. One is low sensitivity (Fig. 4B—B-Raf). This could be due either to low epitope abundance, or poor antibody affinity; in this case we have prior to data estimating low nanomolar concentration of total B-Raf in this cell lysate (MCF10A)[29]. This suggests there is simply not enough epitope to effectively quantify, rather than an issue with the antibody itself. However, we do note that we were unable to validate the antibody against total C-Raf, whereas that of phospho-C-Raf was validated at much lower working concentrations than used for the total C-Raf antibody. This suggests at least for the total C-Raf antibody, there could be issues of affinity hindering quantitative use. A similar trend was observed for GSK3beta (see also above). In other cases (SAPK/JNK), we noted technical issues in the run (Fig. 4C), which required additional evaluation before deeming the antibody valid for quantitative use. Subsequent runs provided data that allowed validation of this antibody. In either case, there potentially seemed not to be an issue with the antibodies themselves, but rather other aspects of the experiment (cell system or technical).

### Comparison to Macro-Western

An antibody may have adequate quantitative performance in microwestern, but it is unclear what that implies for traditional “macro”-western. Therefore, we performed analogous macro-western experiments using two antibodies that were validated via microwestern—phospho-ERK and α-tubulin. Images and resulting quantification are presented in Fig. 5A. Acceptable quantitative behavior is observed for both antibodies via macro-western. We used those same antibodies at the same dilutions in independent microwestern experiments (Fig. 5B). These results showed a similarly valid quantitative behavior. Thus, we conclude that validation results from microwestern are likely to carry over to macro-western. Of course, we have not tested every antibody in such a manner, since microwestern allows much larger throughput than macro-western. However, these results certainly provide confidence in translation to the traditional western blot. This is particularly satisfying given the much smaller amount of protein sample required for microwestern (Fig. 5C). We also note in the microwestern results, a slight tendency towards saturation at the higher sample amounts (Fig. 5B). However, since macro-western seems to have a larger linear range in these cases, one may consider the scaling conservative, as linear range seems to grow with the assay scale, at least with these examples.

**Figure 5.**
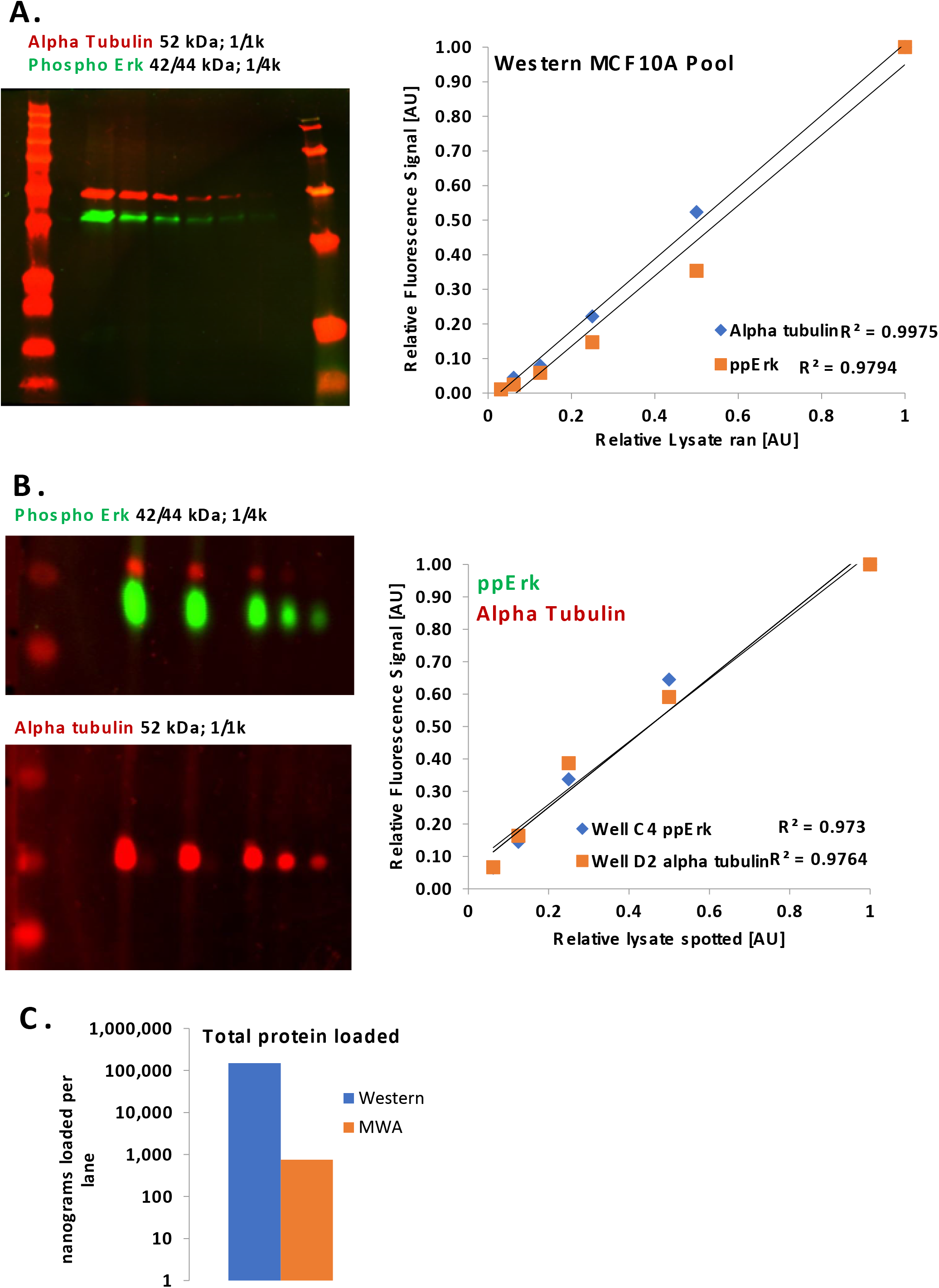
Comparison of Western to Microwestern. Lysates from exponentially growing MCF10A cells were used for these experiments. A. Measurement of ppERK and alpha-tubulin using a “regular” Western with quantification results. B. Measurement of ppERK and alpha-tubulin using a Microwestern at matched antibody dilution conditions with the same cell systems. C. Total protein loaded per well is significantly lower for a MWA. Note the log y-axis scale.

## CONCLUSIONS

A key aspect of reproducibility is the fidelity of research reagents. Antibodies are a workhorse in biomedical science, and their validity is often not investigated and/or is highly dependent on the assay of choice. Here we demonstrate the use of meso-scale western blotting—the microwestern array—to evaluate the suitability of antibodies for quantitative use in western blotting. By providing the results publicly it can be re-used by other researchers for similar purposes to provide confidence in their quantitation.

## ACKNOWLEDGEMENTS

We are grateful to the NIH LINCS program for funding (U54HG008098 to MRB and RI). We acknowledge R01GM104184 (to MRB). AMB and ADS acknowledge an NIGMS-funded Integrated Pharmacological Sciences Training Program grant (T32GM062754) for support.

